# The Pathogenicity of SARS-CoV-2 in hACE2 Transgenic Mice

**DOI:** 10.1101/2020.02.07.939389

**Authors:** Linlin Bao, Wei Deng, Baoying Huang, Hong Gao, Jiangning Liu, Lili Ren, Qiang Wei, Pin Yu, Yanfeng Xu, Feifei Qi, Yajin Qu, Fengdi Li, Qi Lv, Wenling Wang, Jing Xue, Shuran Gong, Mingya Liu, Guanpeng Wang, Shunyi Wang, Zhiqi Song, Linna Zhao, Peipei Liu, Li Zhao, Fei Ye, Huijuan Wang, Weimin Zhou, Na Zhu, Wei Zhen, Haisheng Yu, Xiaojuan Zhang, Li Guo, Lan Chen, Conghui Wang, Ying Wang, Xinming Wang, Yan Xiao, Qiangming Sun, Hongqi Liu, Fanli Zhu, Chunxia Ma, Lingmei Yan, Mengli Yang, Jun Han, Wenbo Xu, Wenjie Tan, Xiaozhong Peng, Qi Jin, Guizhen Wu, Chuan Qin

## Abstract

Severe acute respiratory syndrome CoV-2 (SARS-CoV-2) caused the Corona Virus Disease 2019 (COVID-19) cases in China has become a public health emergency of international concern (PHEIC). Based on angiotensin converting enzyme 2 (ACE2) as cell entry receptor of SARS-CoV, we used the hACE2 transgenic mice infected with SARS-CoV-2 to study the pathogenicity of the virus. Weight loss and virus replication in lung were observed in hACE2 mice infected with SARS-CoV-2. The typical histopathology was interstitial pneumonia with infiltration of significant lymphocytes and monocytes in alveolar interstitium, and accumulation of macrophages in alveolar cavities. Viral antigens were observed in the bronchial epithelial cells, alveolar macrophages and alveolar epithelia. The phenomenon was not found in wild type mice with SARS-CoV-2 infection. The pathogenicity of SARS-CoV-2 in hACE2 mice was clarified and the Koch’s postulates were fulfilled as well, and the mouse model may facilitate the development of therapeutics and vaccines against SARS-CoV-2.

---

In late December of 2019, the coronavirus disease 2019 (COVID-19) caused by severe acute respiratory syndrome CoV-2 (SARS-CoV-2), linked to a seafood market in which exotic animals were also sold and consumed, were identified and reported from Wuhan City, Hubei Province, China^1,2^. The number of confirmed cases has since soared, with almost 78,000 cases reported and over 2,700 deaths as of February 25, 2020 in China^3^, and imported cases from travelers of mainland China in several other countries. It is critical to find the pathogenicity and biology of the virus for prevention and treatment of the disease.

Because SARS-CoV-2 was highly homologous with severe acute respiratory syndrome coronavirus (SARS-CoV), human angiotensin-converting enzyme 2 (hACE2), which was the entry receptor of SARS-CoV, was also considered to have a high binding ability with the SARS-CoV-2 by molecular biological analysis^4,5^. Therefore, we used the hACE2 transgenic and wild type (WT) mice infected with SARS-CoV-2 to study the pathogenicity of the virus.

Specific pathogen-free, 6-11-month-old, male and female WT mice (WT-HB-01, n=15) and hACE2 mice (ACE2-HB-01, n=19) were inoculated intranasally with SARS-CoV-2 stock virus (HB-01) at a dosage of 10^5^ TCID_50_/50 μl inoculum volume per mouse after the mice were anesthetized by 2.5% avertin, and the mock-treated hACE2 mice (ACE2-Mock, n=15) were used as control. Clinical manifestations were recorded from thirteen mice (WT-HB-01, n=3; ACE2-Mock, n=3; ACE2-HB-01, n=7). Compared to WT-HB-01 mice and ACE2-Mock mice, slight bristles and weight loss were only observed in ACE2-HB-01 mice during the 14 days observation, and other clinical symptoms such as arched back and decreased projection of external stimuli were not found. Notably, the weight loss of ACE2-HB-01 mice was up to 8% at 5 days post infection (dpi) (Figure 1a).

**Figure 1.**
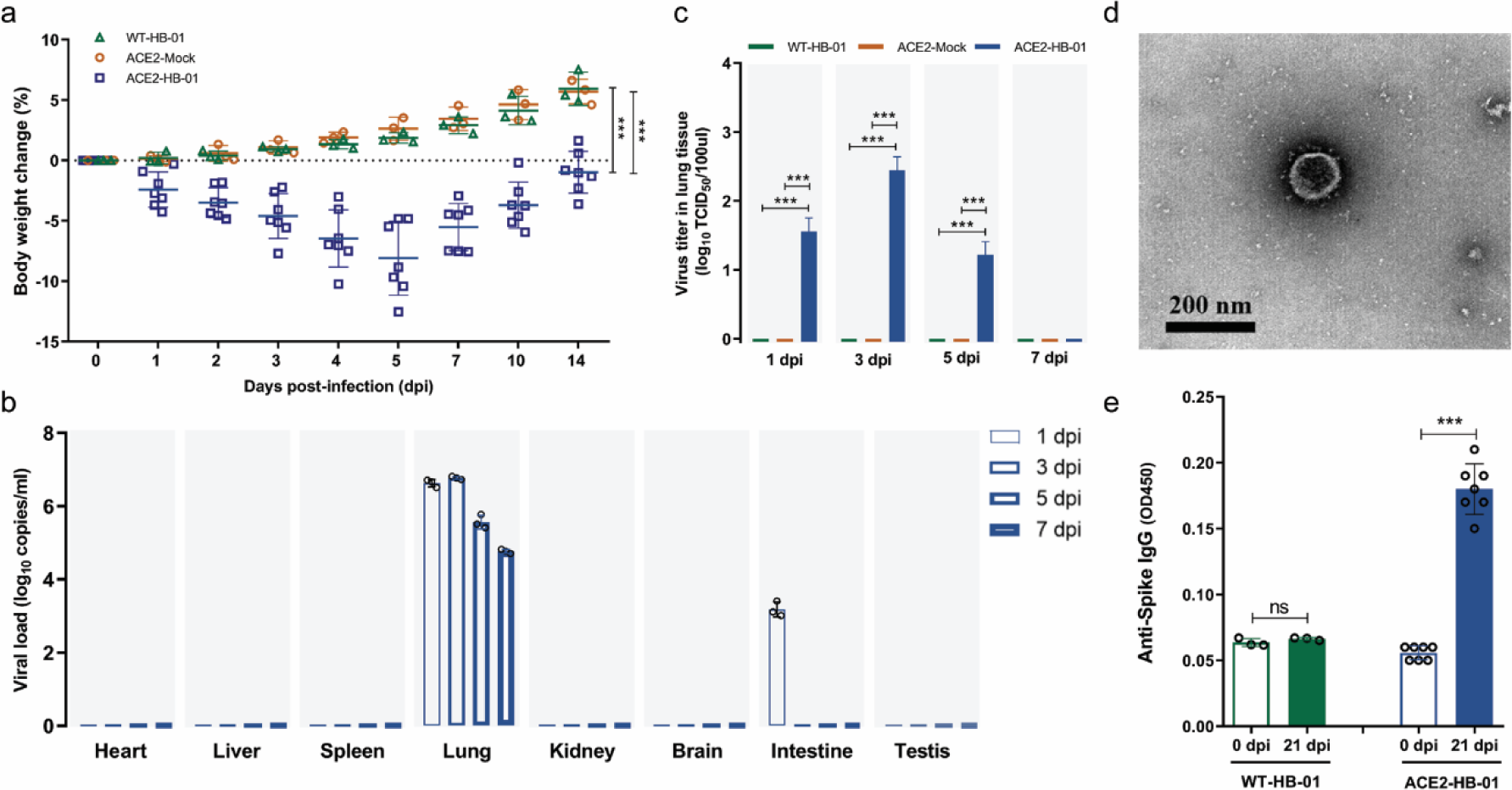
Weight loss, virus replication and specific IgG production in hACE2 mice post infection with SARS-CoV-2. For weigh loss record, hACE2 mice (n=7) and WT mice (n=3) were experimentally challenged intranasally with SARS-CoV-2, and the ACE2-Mock mice (n=3) were used as control. And then the weight loss was recorded over 14 days (a, ANOVA, \*\*\**p*<*0.001*). To screen virus replication, 12 mice were infected in each group, and 3 mice per group were sacrificed and their major organs harvested for viral load and virus titer at 1 dpi, 3 dpi, 5 dpi and 7 dpi respectively. The distribution of SARS-CoV-2 in the primary organs of ACE2-HB-01 mice was detected by qRT-PCR (b). Virus titers of lungs were determined on Vero E6 cells (c, unpaired *t*-test, \*\*\**p*<*0.001*), and the virus isolated from lungs of ACE2-HB-01 mice at 3 dpi was observed by electron microscope (d). The specific IgG against SARS-CoV-2 was detected at 21 dpi by ELISA (e, unpaired *t*-test, \*\*\**p*<*0.001*).

Next, viral replication and pathological changes were examined from three animals per group at each time point, and the primary organs were collected periodically, including heart, liver, spleen, lung, kidney, brain, intestine and testis. As shown in Figure 1b, viral loads were detected by qRT-PCR at 1 dpi, 3 dpi, 5 dpi and 7 dpi in the lungs of ACE2-HB-01 mice but not in that of WT-HB-01mice (data not shown), and the viral RNA copies reached a peak at 3 dpi (10^6.77^ copies/ml). Interestingly, the viral RNA was also detected at 1 dpi in the intestine of ACE2-HB-01 mice, which was failed to be detected in other tissues along the timeline (Figure 1b). Consistent to the results of viral loads, infectious virus was respectively isolated from the lungs of ACE2-HB-01 mice at 1 dpi, 3 dpi and 5 dpi, and the highest virus titers were detected at 3 dpi (10^2.44^ TCID_50_/100 μl) (Figure 1c). Meanwhile, the infectious virus was isolated by Vero E6 cells culture from lung, and the SARS-CoV-2 particles were observed by electron microscopy (Figure 1d). However, the virus was failed to be isolated from the lungs in WT-HB-01 mice and ACE2-Mock mice along the detecting timeline (Figure 1c), which suggested that hACE2 was essential for SARS-CoV-2 infection and replication in mice. Moreover, specific IgG antibodies against S protein of SARS-CoV-2 were positively measured in the sera of ACE2-HB-01 mice at 21 dpi (Figure 1e).

There were no obviously gross and histopathological changes at 1 dpi in all the animals of each group. Compared to WT-HB-01 mice or ACE2-mock mice with homogeneously pink and slightly deflated, all the ACE2-HB-01 mice at 3 dpi displayed comparable gross lesions with focal to multifocal dark red discoloration in certain lung lobes. The lesions progressed into multifocal to coalescent scattered-dark reddish purple areas and focal palpable nodules throughout the lung lobes at 5 dpi (Figure 2a). The damaged lungs became swollen and enlarged in size. Microscopically, consistently, lung tissues from ACE2-HB-01 mice at 3 dpi showed moderate, multifocal, interstitial pneumonia. Inflammatory cells including lymphocytes and monocytes accumulated in the alveolar interstitium and caused thickening of the alveolar walls (Figure 2a). Increased collagen fiber in the thickened alveolar interstitium in the ACE2-HB-01 mice was confirmed by Modified Masson’s Trichrome stain with blue colors (Supplementary Figure 1a). The bronchiolar epithelial cells showed swelling, degeneration, some of which dissolved (Figure 2a). Affected bronchioles were filled with occasionally small amounts of periodic acid-Schiff (PAS) positive-exudation or denatured and detached bronchiolar epithelium (Supplementary Figure 1b). The alveolar cavities were dispersed mainly by swollen and degenerative alveolar macrophages and lymphocytes (Figure 2a). To investigate the infiltration of specific inflammatory cells, immunohistochemical (IHC) was carried out to identify MAC2^+^ macrophages (Supplementary Figure 2a), CD3^+^ T lymphocytes, and CD19^+^ B lymphocytes (Supplementary Figure 2b). Compared to the WT-HB-01 mice, more macrophages, T lymphocytes, and B lymphocytes were observed in the lungs of ACE2-HB-01 mice along with lasting prolonged infection time. The MAC2^+^ macrophages were diffusely infiltrated in the alveolar cavities (at 3 dpi) or focally aggregated together in the thickened alveolar septum (at 5 dpi, Supplementary Figure 2a). CD3^+^ T lymphocytes and CD19^+^ B lymphocytes were scattered dispersed or occasionally aggregated in the alveolar interstitium in the ACE2-HB-01 mice, by contrast, they were rarely observed in the WT-HB-01 mice (Supplementary Figure 2b). Perivascular infiltrated inflammatory cells, including lymphocytes and monocytes, were observed multifocally within and adjacent to affected areas of the lungs. At 5 dpi, the lung progressed into coalescing interstitial pneumonia with diffuse lesions. Thickened alveolar septa were filled with lymphocytes and monocytes (Figure 2a). IHC staining of sequential section revealed that viral antigens were found in the alveolar macrophages, alveolar epithelia, and the degenerative and being desquamated bronchial epithelial cells (Figure 2b). There were no significant histopathological lesions (Supplementary Figure 3) or viral antigens of SARS-CoV-2 (Supplementary Figure 4) in the other organs, including myocardium, liver, spleen, kidney, cerebrum, intestine, and testis. At 7 dpi, the pneumonia became mild with focal lesion areas (data not shown).

**Figure 2.**
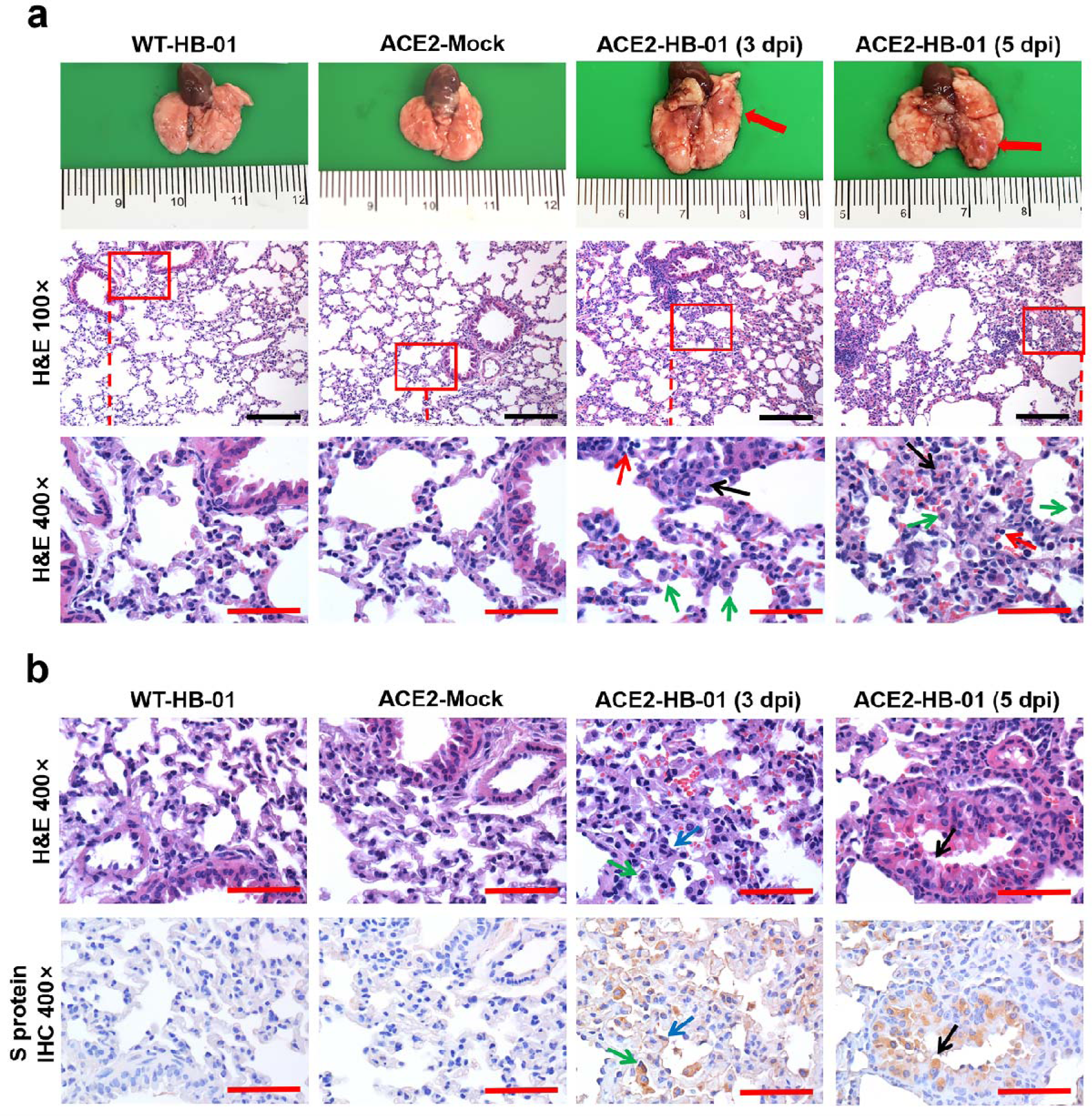
Gross pathology, histopathology, and immunohistochemistry of the lungs in SARS-CoV-2-infected hACE2 mice. a. Gross pathology and histopathology of lungs from WT-HB-01 mice (3 dpi), ACE2-Mock mice (3 dpi) and ACE2-HB-01 mice (3 dpi and 5 dpi). Postmortem examinations showed focal dark red lesions (red arrow) throughout the dorsal of the right middle lobe of the lung at 3dpi. The lesions progressed into multifocally scattered-dark reddish purple areas and palpable nodules (red arrow) throughout the right lobe of the lung at 5 dpi. Histopathological observation indicated that moderate interstitial pneumonia with thickened alveolar septa (black arrows) and infiltration of lymphocytes (red arrows). The swollen and degenerative alveolar macrophages (green arrows) are scattered within the alveolar cavities at 3dpi and 5 dpi. b. Immunohistochemical examination of lungs of each group. The sequential sections were stained by HE and IHC, respectively. The viral antigens were observed in the alveolar macrophage (green arrows), the alveolar epithelia (blue arrows), and the degenerative and being desquamated bronchial epithelial cells (black arrows). Black bar = 100 µm, red bar = 50 µm.

In addition, the co-localization of SARS-CoV-2 S protein (Fifure 3f) and hACE2 receptor (Figure 3g) was demonstrated in alveolar epithelial cells of ACE2-HB-01 mice by immunofluorescence (Figures 3h). The phenomenon was not found in the ACE2-Mock mice (Figures 3a, b, c and d) or WT-HB-01 mice (data not shown), indicating that the SARS-CoV-2, as same as SARS-CoV, also utilizes the hACE2 as a receptor for entry^4^.

**Figure 3.**
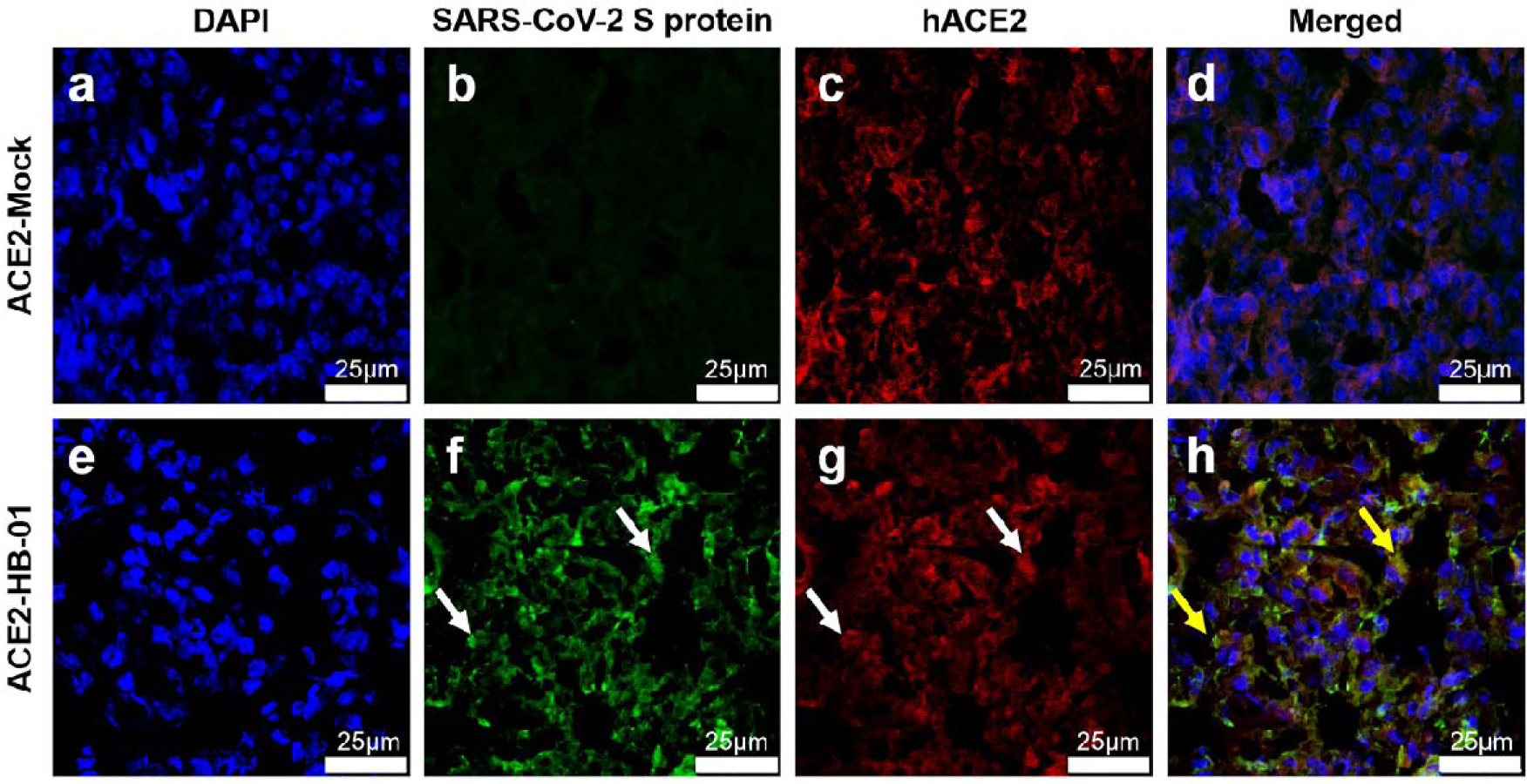
Immunofluorescence analysis of viral antigens in lungs of SARS-CoV-2-infected hACE2 mice. Co-localization of SARS-CoV-2 S protein and hACE2 receptor in hACE2 mouse lungs, the sections were incubated with anti-SARS-CoV-2 S protein antibody, anti-human ACE2 antibody, and DAPI. The lung sections of ACE2-Mock mice (a-d). The lung sections of ACE2-HB-01 mice (e-h). The white arrows showed the viral S protein (f) and hACE2 (g), respectively, the yellow arrow showed the merge of viral S protein and hACE2 (h). White bar=25 µm.

The speed of geographical spread of COVID-19 caused by SARS-CoV-2 has been declared as public health emergency of international concern (PHEIC), with cases reported on multiple continents only weeks after the disease was first reported^6^. Although it has been determined by bioinformatics that the pathogen of this epidemic is a novel coronavirus, it is necessary to be confirmed by animal experiments following Koch’s postulates.

In the present study, after the experimental infection of hACE2 transgenic mice with one of the earliest known isolates of SARS-CoV-2, the mice lost weight and showed interstitial pneumonia, which are comparable with initial clinical reports of pneumonia caused by SARS-CoV-2^7^. Meanwhile, virus replication was detected in the lung of ACE2-HB-01 mice and elicited the specific IgG against SARS-CoV-2. In clinical autopsy, histological examination showed bilateral diffuse alveolar damage with cellular fibromyxoid exudates^8^. Correspondingly, ACE2-HB-01 mice showed coalescing interstitial pneumonia in this study. Furthermore, specifically histopathology changes and progress were observed in the mice, e.g. affected bronchioles were filled with occasionally small amounts of periodic acid schiff (PAS) positive-exudation or denatured and detached brochiolar epithelium.

The case fatality rate of currently reported cases is about 2%, which implies that so far, this novel coronavirus does not seem to cause the high fatality rates as SARS-CoV (9-11%)^9^, which suggested the difference in pathogenicity between the two viruses. The pathogenicity of SARS-CoV-2 seems mild compared to SARS-CoV in mice, the latter caused extrapulmonary organ damage, includes brain, kidney, intestine, heart and liver, furthermore, the neurons are susceptible for SARS-CoV infection, and cerebral casculitis and hemorrhage were observed in hACE2 transgenic mice^10,11^. However, only interstitial pneumonia was observed in SARS-CoV-2-infected hACE2 mice, implying the disparity in pathogenicity of the coronavirus.

Therefore, our results clarified the pathogenicity of SARS-CoV-2 in mice, together with the previous clinical studies^7^, completely fulfills the Koch’s postulates^12^ and confirmed SARS-CoV-2 was the pathogen of COVID-19. The mouse model may facilitate the development of therapeutics and vaccines against SARS-CoV-2.

## Materials and methods

### Ethics statement

Murine studies were performed in an animal biosafety level 3 (ABSL3) facility using HEPA-filtered isolators. All procedures in this study involving animals were reviewed and approved by the Institutional Animal Care and Use Committee of the Institute of Laboratory Animal Science, Peking Union Medical College (BLL20001).

### Viruses and cells

The SARS-CoV-2 (strain HB-01) was kindly provided by Professor Wenjie Tan^1^, from the China Centers for Disease Control and Prevention (China CDC). The complete genome for this SARS-CoV-2 was submitted to GISAID (BetaCoV/Wuhan/IVDC-HB-01/2020|EPI_ISL_402119), and deposited in the China National Microbiological Data Center (accession number NMDC10013001 and genome accession numbers MDC60013002-01). Seed SARS-CoV-2 stocks and virus isolation studies were performed in Vero cells, which are maintained in Dulbecco’s modified Eagle’s medium (DMEM, Invitrogen, Carlsbad, USA) supplemented with 10% fetal bovine serum (FBS), 100 IU/ml penicillin, and 100 µg/ml streptomycin, and incubated at 37°C, 5% CO_2_. For infected mice, the lung homogenates were used for virus titration tests using endpoint titration in Vero E6 cells. Virus titer of the supernatant were determined using a standard 50% tissue culture infection dose (TCID50) assay.

### Animal experiments

For the animal experiments, specific pathogen-free, 6-11-month-old, male and female transgenic hACE2 mice were obtained from the Institute of Laboratory Animal Science, Peking Union Medical College, China. Transgenic mice were generated by microinjection of the mice ACE2 promoter driving the human ACE2 coding sequence into the pronuclei of fertilized ova from ICR mice, and then human ACE2 integrated was identified by PCR as previous described^10^, the hACE2 mainly expressed in lung, heart, kidney, and intestine of transgenic mice. The hACE2 mice or WT (ICR) mice were respectively inoculated intranasally with SARS-CoV-2 stock virus at a dosage of 10^5^ TCID_50_ and hACE2 mice intranasally inoculated with equal volume of PBS was used as mock-infection control. The infected animals were continuously observed daily to record body weights, clinical symptoms, responsiveness to external stimuli and death. And the mice were dissected at 1 dpi 3 dpi, 5 dpi and 7 dpi respectively to collect different tissues to screen virus replication and histopathological changes.

### Preparation of Homogenate Supernatant

Tissues homogenates were prepared by homogenizing perfused whole tissue using an electric homogenizer for 2 min 30 s in 1 ml of DMEM. The homogenates were centrifuged at 3,000 rpm for 10 min at 4°C. The supernatant was collected and stored at −80°C for viral isolation and viral load detection.

### RNA extraction and qRT-PCR

Total RNA was extracted from organs using the RNeasy Mini Kit (Qiagen, Hilden, Germany), and reverse transcription was performed using the PrimerScript RT Reagent Kit (TaKaRa, Japan) following manufacturer instructions. A quantitative real-time reverse transcription-PCR (qRT-PCR) reactions were performed using the PowerUp SYBG Green Master Mix Kit (Applied Biosystems, USA), in which samples were processed in duplicate using the following cycling protocol: 50°C for 2 min, 95°C for 2 min, followed by 40 cycles at 95°C for 15 s and 60°C for 30 s, and then 95°C for 15 s, 60°C for 1 min, 95°C for 45 s. The primer sequences used for RT-PCR are targeted against the envelope (E) gene of SARS-CoV-2 and are as follows: Forward: 5’-TCGTTTCGGAAGAGACAGGT-3’, Reverse: 5’-GCGCAGTAAGGATGGCTAGT-3’. The PCR products were verified by sequencing using the dideoxy method on an ABI 3730 DNA sequencer (Applied Biosystems, CA, USA). During the sequencing process, amplification was performed using specific primers. The sequences for this process are available upon request. The sequencing reads obtained were linked using DNAMAN, and the results were compared using the Megalign module in the DNAStar software package.

### ELISA method

The specific IgG against SARS-CoV from ACE2-HB-01 mice and WT-HB-01 mice were determined by enzyme-linked immunosorbent assay (ELISA). 96-well plates were coated with Spike 1 protein of SARS-CoV-2 (0.1μg/100 μL, Sino Biological, 40591-V08H), the tested sera were diluted at 1:100 and added to each well, and 3 multiple wells were set for each sample, and then incubated at 37°C for 30 minutes, followed by the goat anti-mouse secondary antibodies conjugated with HRP (ZB-2305, zhongshan,1:10,000 dilution), and incubated at room temperature for 30 minutes. The reaction was developed by TMB substrate and the optical densities at 450 nm were determined (Metertech960 enzyme marker with 450 nm wavelength).

### Laboratory preparation of the antibody of SARS-CoV-2 Spike-1 (S1) protein

Mice were immunized with purified SARS-CoV-2 S1 protein (Sino biological) and splenocytes of hyper immunized mice were fused with myeloma cells. Positive clones were selected by ELISA using SARS-CoV-2 S1 protein (Supplementary Figure 1). The cell supernatant of 7D2 clone, binding to SARS-CoV-2 S1 protein, was collected for immunofluorescence analysis.

### Pathological Examination

Autopsies were performed in the animal biosafety level 3 (ABSL3) laboratory. Major organs were grossly examined and then fixed in 10% buffered formalin solution, and paraffin sections (3-4 µm in thickness) were prepared routinely. Hematoxylin and Eosin (H&E) stain, periodic acid-Schiff (PAS) stain and modified Masson’s Trichrome stain were used to identify histopathological changes in all the organs. The histopathology of the lung tissue was observed by light microscopy.

### Immunohistochemistry (IHC)

The organs were fixed in 10% buffered formalin solution, and paraffin sections (3-4 µm in thickness) were prepared routinely. Sections were treated with an antigen retrieval kit (Boster, AR0022) for 1 min at 37°C and quenched for endogenous peroxidases in 3% H_2_O_2_ in methanol for 10 min. After blocking in 1% normal goat serum, the sections were incubated with 7D2 monoclonal antibody (laboratory preparation) at 4 °C overnight, followed by HRP-labeled goat anti-mouse IgG secondary antibody (HRP) (Beijing ZSGB Biotechnology, ZDR-5307). Alternatively, the sections were stained with MAC2 antibody (Cedarlane Laboratories, CL8942AP), CD3 antibody (Dako, A0452) or CD19 antibody (Cell Signaling Technology, 3574) at 4°C overnight. Subsequently, the sections were goat anti-rat IgG secondary antibody (HRP) (Beijing ZSGB Biotechnology, PV9004), goat anti-rabbit IgG secondary antibody (HRP) (Beijing ZSGB Biotechnology, PV9001) for 60 min, and visualized by incubation with 3,30-diaminobenzidine tetrahydrochloride (DAB). The slices were counterstained with hematoxylin, dehydrated and mounted on a slide and viewed under an Olympus microscope.

### Confocal microscopy

For viruses and hACE2 receptor co-localization analysis, the lung tissue sections were washed twice with PBS, fixed by Immunol Staining Fix Solution (P0098), blocked 1 hour at room temperature by Immunol Staining Blocking Buffer (P0102) and then incubated overnight at 4°C with the appropriate primary and secondary antibodies. The nuclei were stained with DAPI. Anti-S protein antibody (mouse monoclonal 7D2, laboratory preparation, 1:200) and anti-hACE2 antibody (rabbit polyclonal, ab15348, Abcam1:200) were used as the primary antibody. The sections were washed with PBS and incubated with secondary antibodies conjugated with FITC (goat anti-mouse, ZF-0312, Beijing ZSGB Biotechnology, 1:200) and TRITC (goat anti rabbit, ZF-0317, Beijing ZSGB Biotechnology, 1:200), dried at room temperature and observed via fluorescence microscopy. For the expression of hACE2, the sections from WT mice stained with anti-ACE2 antibody were used as the negative control, and the stable cell line expressing hACE2 was used as the positive control. For the viral antigen, the sections from ACE2-Mock mice incubated with anti-S protein antibody were used as the negative control.

### Transmission Electron Microscopy

Supernatant from Vero E6 cell cultures that showed cytopathic effects was collected, inactivated with 2% paraformaldehyde for at least 2 hours, and ultracentrifuged to sediment virus particles. The enriched supernatant was negatively stained on film-coated grids for examination. The negative stained grids were observed under transmission electron microscopy.

### Statistical analysis

All data were analyzed with GraphPad Prism 8.0 software. Statistically significant differences between two groups were determined using unpaired Student’s *t*-tests. The statistical significance among three groups was assessed by one-way ANOVA. A two-sided p value*<0.05* was considered statistically significant.

## Acknowledgement

We thank Dr George F Gao for his advice and coordination on this work. We thank Hongkui Deng, Xiuhong Yang and Lianfeng Zhang for providing the hACE2 mice as a gift. We also thank Gary Wong for helping us proofread the language. This work was supported by National Research and Development Project of China (Grant No. 2020YFC0841100), Fundamental Research Funds for CAMS of China (2020HY320001), National Key Research and Development Project of China (Grant No. 2016YFD0500304), CAMS initiative for Innovative Medicine of China (Grant No. 2016-I2M-2-006), and National Mega projects of China for Major Infectious Diseases (2017ZX10304402).

---

**Supplementary Figure 1.**
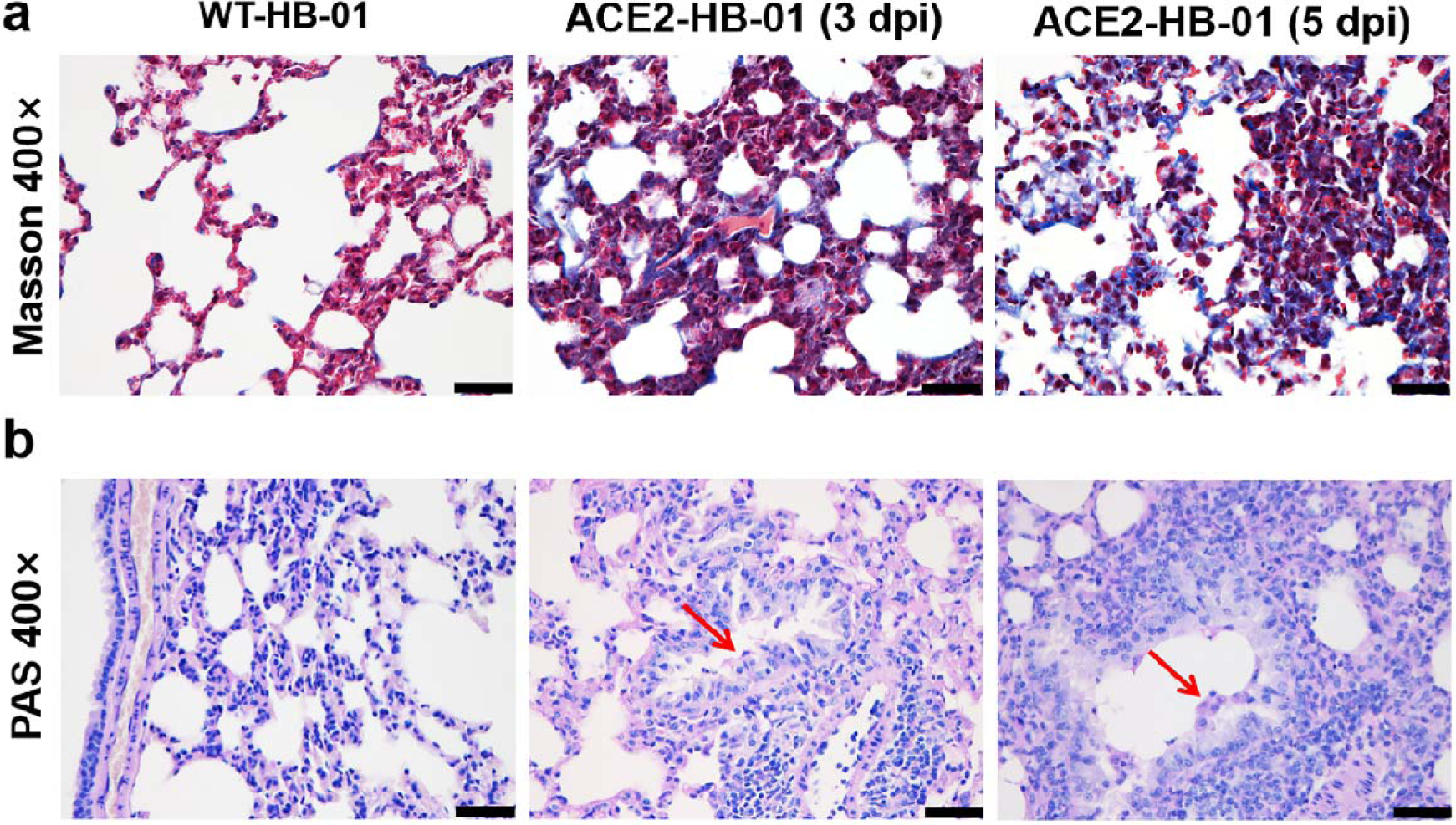
Special stains of the lungs in the WT-HB-01 and ACE2-HB-01 mice at 3dpi and 5 dpi. a. Modified Masson’s Trichrome of the lung. Compared to the WT-HB-01 group, the increased collagen fiber (blue-stained fibers) in the thickened alveolar interstitium were observed in both the ACE2-HB-01 mice at 3dpi and 5 dpi. b. Periodic acid schiff (PAS) staining of the respiratory epithelium in bronchioles. A small amount of mucus was accumulated on the surface of bronchial epithelial cells. Black bar=40 µm.

**Supplementary Figure 2.**
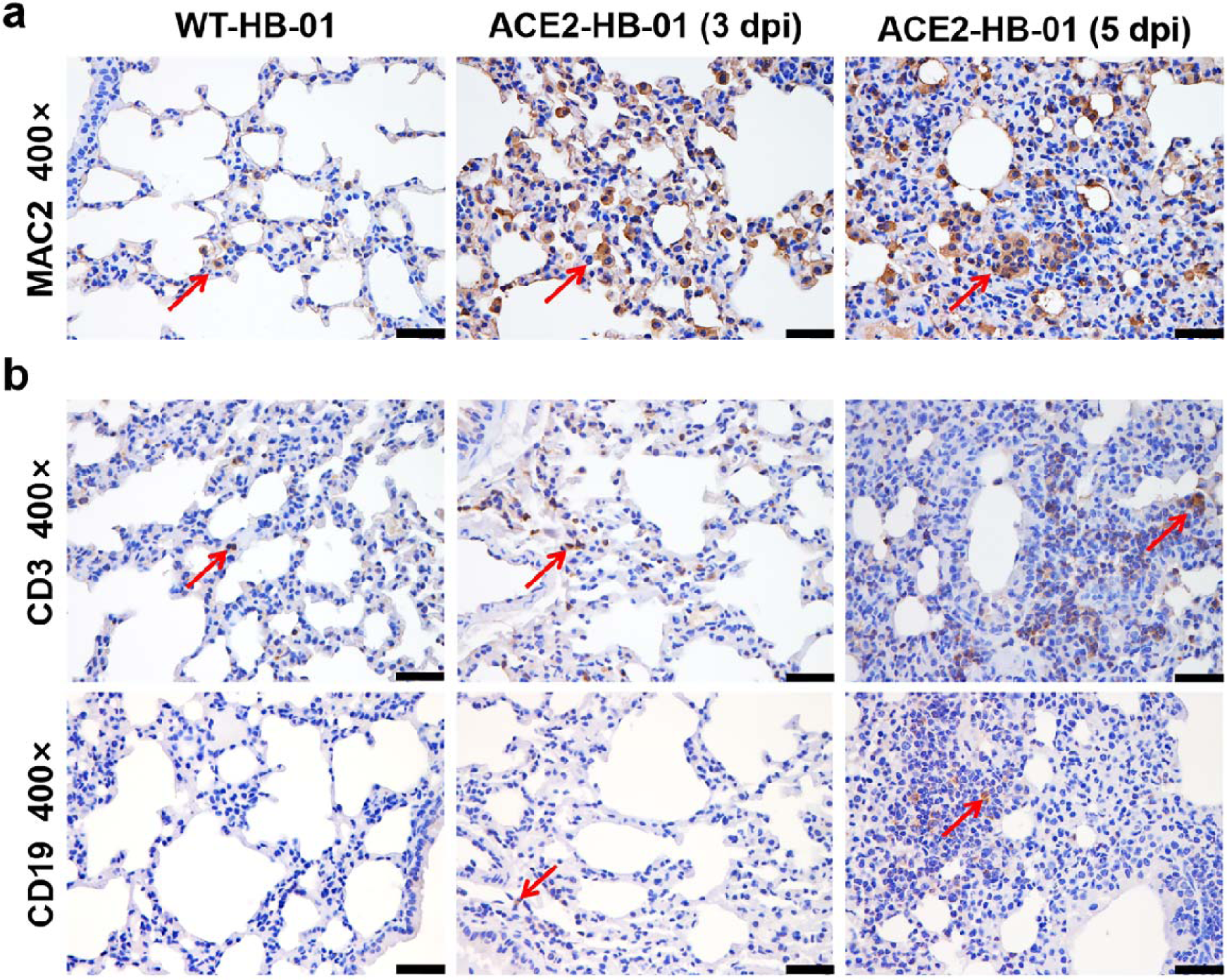
IHC was carried out to identify macrophages with MAC2, T lymphocytes with CD3, and B lymphocytes with CD19. a. Diffuse infiltration of macrophages (red arrow) in the expanded alveolar septum in the ACE2-HB-01 mice at 3 dpi and 5 dpi. b. Many T lymphocytes (red arrow) infiltrated the thickened alveolar septa in the first row of b at 5 dpi in the ACE2-HB-01 mice. A few B lymphocytes (red arrow) were observed in the ACE2-HB-01 mice. Black bar=40 µm.

**Supplementary Figure 3.**
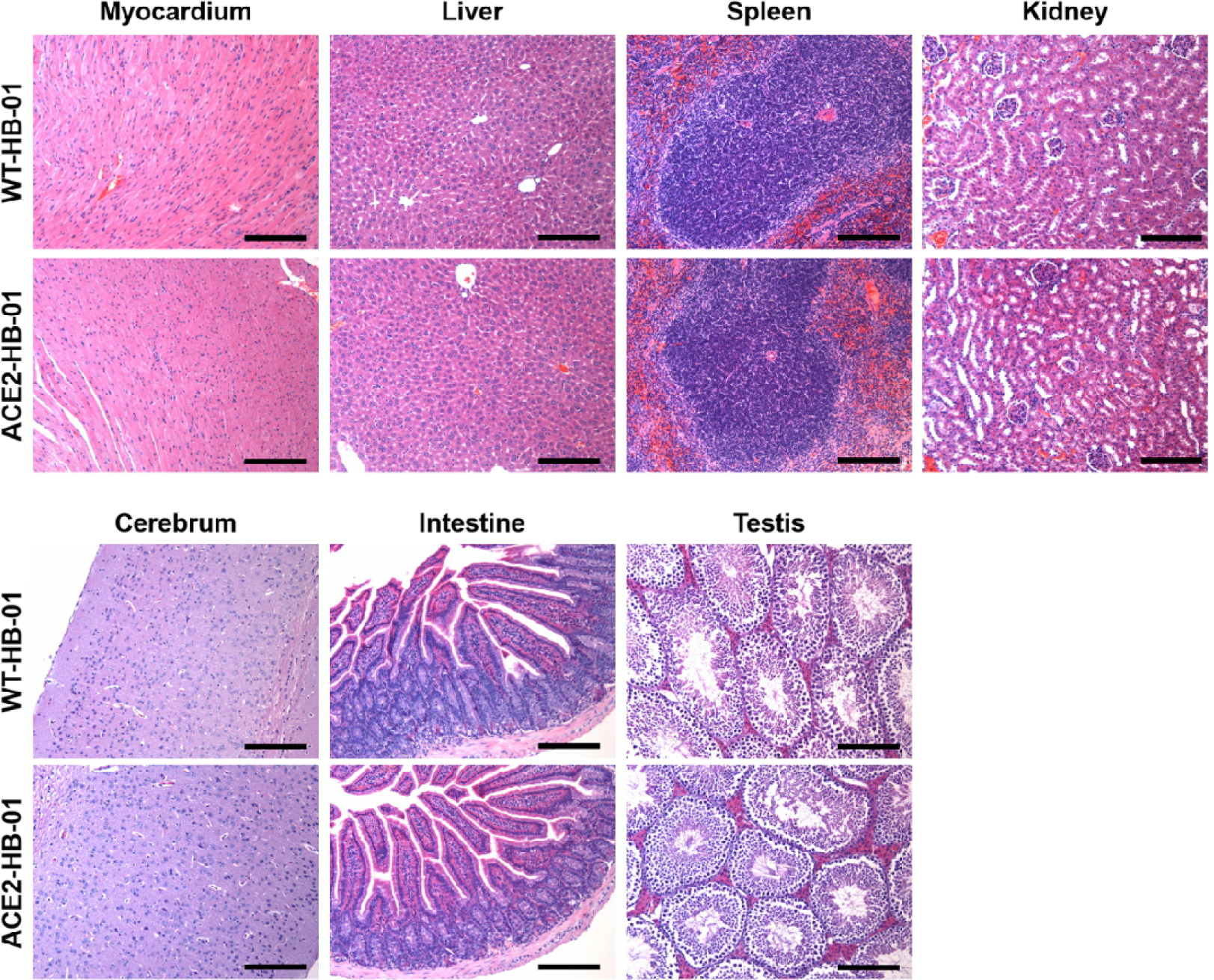
The histopathological observation of the organs in the WT-HB-01 and ACE2-HB-01 mice. There were no significant histopathological lesions in the organs, including myocardium, liver, spleen, kidney, cerebrum, intestine and testis in the ACE2-HB-01 mice compared with the ACE2-Mock mice. Black bar = 100 µm.

**Supplementary Figure 4.**
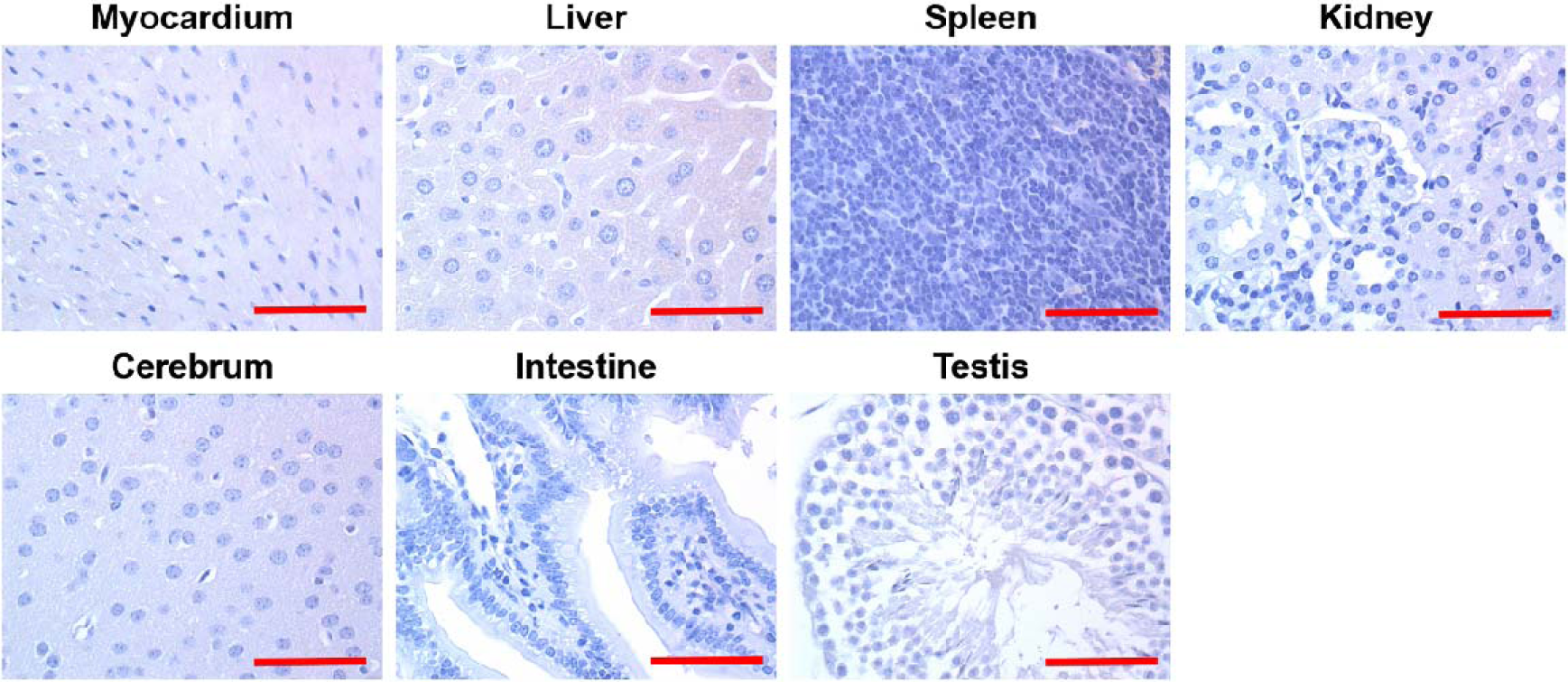
The immunohistochemical (IHC) observation of the organs in the ACE2-HB-01 mice. There were no SARS-CoV-2 antigens in the organs, including myocardium, liver, spleen, kidney, cerebrum, intestine and testis. Red bar = 50 µm.

**Supplementary Figure 5.**
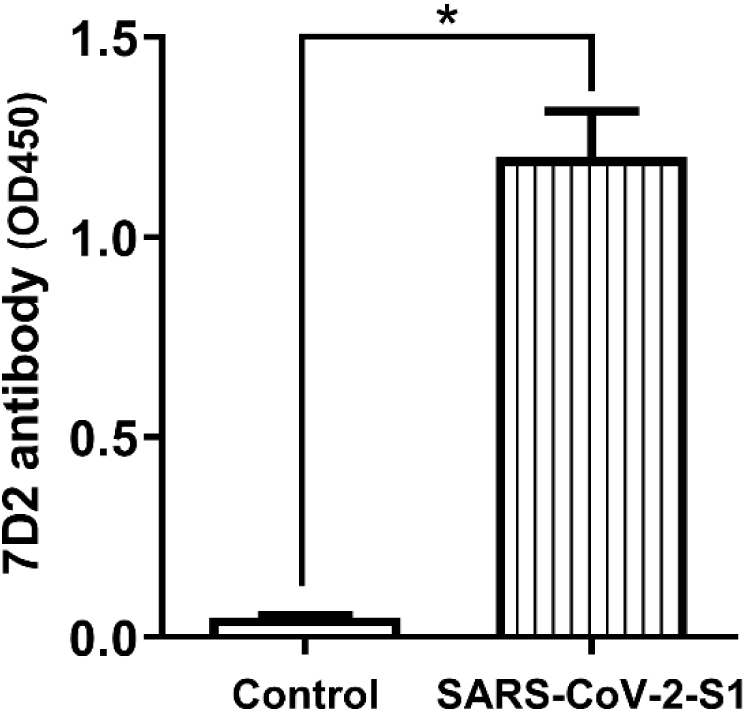
Identification of 7D2 antibody against SARS-CoV-2 S1 protein. The plate coated by 0.2 ug SARS-CoV-2 S1 protein was incubated with 7D2 antibody as primary antibody (1:200) and detected using HRP-conjugated goat anti-mouse secondary antibody. The titer of antibody was determined using enzyme-linked immunosorbent assay (ELISA) assay. Significant differences are indicated with an asterisk (unpaired *t*-test, \**p*<*0.05*).

